# Synaptic vesicle glycoprotein 2 enables viable aneuploidy following centrosome amplification

**DOI:** 10.1101/2025.02.19.639165

**Authors:** Jane E. Blackmer, Erin A. Jezuit, Archan Chakraborty, Ruth A. Montague, Nora G. Peterson, William Outlaw, Donald T. Fox

## Abstract

Amplified centrosome number causes genomic instability, most severely through division into more than two aneuploid daughter cells (multipolar mitosis). Several mechanisms that suppress multipolar division have been uncovered, yet mechanisms that favor viable multipolar division are poorly understood. To uncover factors that promote viability in cells with frequent centrosome amplification and multipolar division, we conducted an unbiased *Drosophila* genetic screen. In 642 mutagenized lines, we exploited the ability of intestinal papillar cells to form and function despite multipolar divisions. Our top hit is an unnamed gene, *CG3168*. We name this gene *synaptic vesicle glycoprotein 2*, reflecting homology to human Synaptic Vesicle Glycoprotein 2 (SV2) proteins. GFP-tagged SV2 localizes to the plasma membrane. In cells with amplified centrosomes, SV2 positions membrane-adjacent centrosomes, which prevents severe errors in chromosome alignment and segregation. Our results uncover membrane-based multipolar division regulation and reveal a novel vulnerability in cells with common cancer properties.

## Introduction

Centrosomes are the microtubule organizing centers (MTOC) of animal cells (Gönczy, 2012, Nigg & Holland, 2018). Duplication of centrosomes is tied to the cell cycle (Banterle & Gönczy, 2017; Fırat-Karalar & Stearns, 2014), and this coordination enables most mitotic cells to exhibit two centrosomes and a bipolar mitotic spindle. Centrosome number amplification (CA) is the property of having greater than two centrosomes per cell (Kiermaier et al., 2024). CA can occur normally, in both mitotic and non-mitotic cells. However, aberrant CA is also a defining property of most tumor cells. In any cell with CA, a multipolar spindle can form (Kalatova et al., 2015; Lingle et al., 2005; Marthiens et al., 2012). These structures present unique challenges to successful mitosis, as chromosomal aneuploidies (imbalances in numbers of each chromosome) can frequently result (Ganem et al., 2009; Silkworth & Cimini, 2012). Aneuploid daughter cells produced from a cell with a multipolar spindle are often missing critical genes and their products, and these cells are frequently inviable (Brinkley, 2001; Vitale et al., 2011). In rarer cases, as was postulated in classic work, CA/multipolar spindles can precede tumor formation (Basto et al., 2008; Boveri, 1902, 2008; Calkins, 1914; Holland & Cleveland, 2009; Levine et al., 2017; LoMastro & Holland, 2019; Nigg, 2002, 2006).

After over a century of study, most of the recent focus on CA has centered on the clustering of extra centrosomes to two poles (Kwon et al., 2008; Marotta et al., 2025; Mercadante et al., 2023; Rhys et al., 2018; Yim et al., 2024). This clustering allows for a bipolar division to occur (**Figure 1A**, **outcome I**). When extra centrosomes cluster to a bipolar spindle, the resulting bipolar anaphase can exhibit erroneous chromosome segregation due to merotelic spindle/kinetochore attachments (**Figure 1A**, **outcome I**) (Ganem et al., 2009; Silkworth & Cimini, 2012). Many conserved centrosome clustering mechanisms have been uncovered, revealing important roles for mitotic regulators such as the Spindle Assembly Complex (SAC), the Chromosomal Passenger Complex (CPC), as well as factors involved in the cytoskeleton and cell adhesion (Kwon et al., 2008; Leber et al., 2010). Thus far, targeting centrosome clustering has only been moderately effective in killing cancer cells with CA (Rebacz et al., 2007; Telentschak et al., 2015).

**Figure 1.**
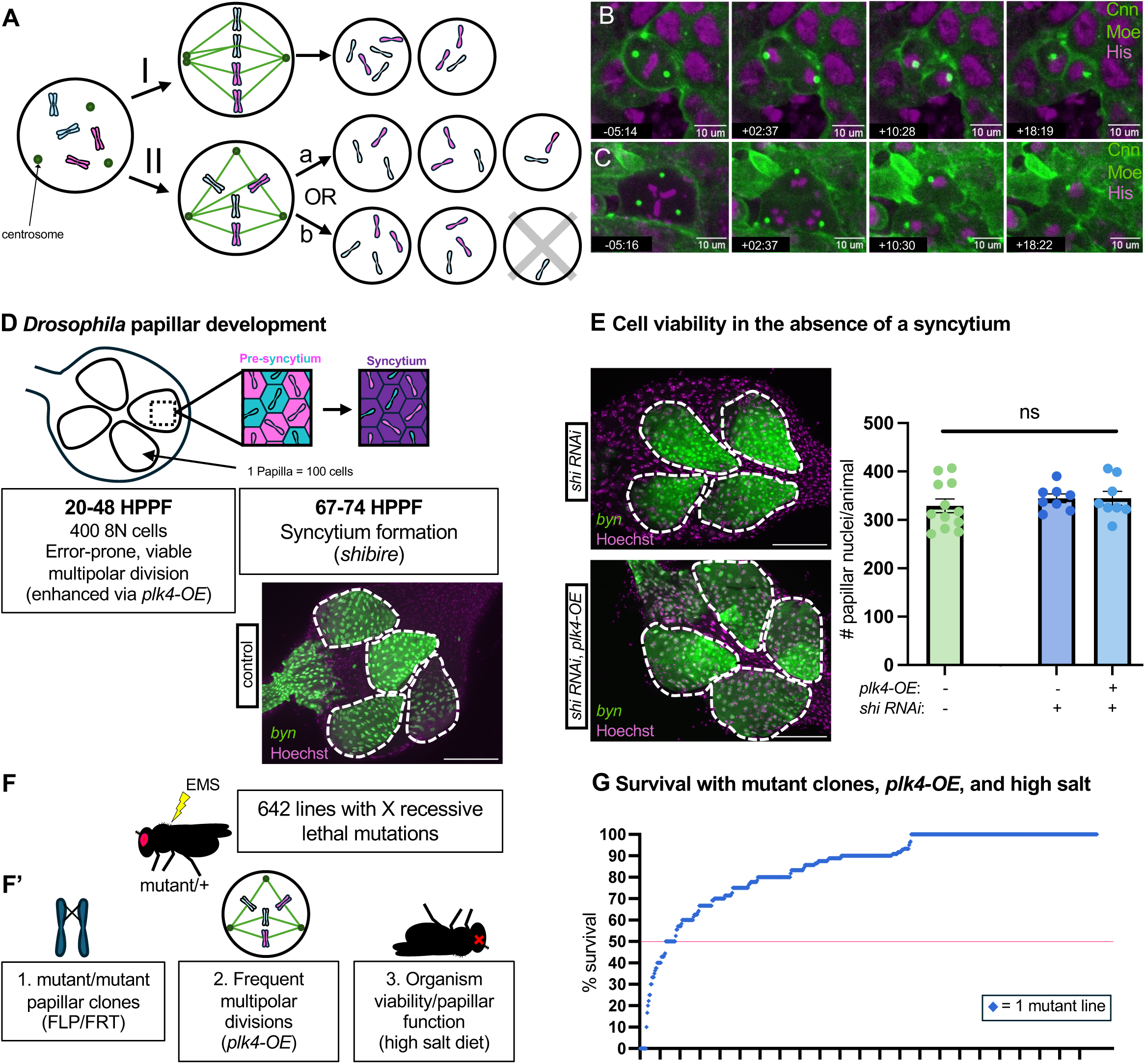
A forward genetic screen for multipolar division survival factors. **(A)** Possible mitotic outcomes of a cell with centrosome number amplification: clustering of centrosomes to form a bipolar spindle and bipolar division (I), Tripolar division with either relatively even distribution of chromosomes to daughter cells allowing for survival (IIa) or with uneven distribution of chromosomes to daughter cells leading to intolerable aneuploidy and cell death in at least one daughter cell (IIb). (**B,C**) *Histone H2AV (His) RFP*, *UAS-centrosomin* (*Cnn)-GFP*;; *brachyenteron*-(*byn)-Gal4, Gal80^ts^*, *UAS-moesin* (Moe)-*GFP* expressing pupal stage wildtype rectal papillar cells undergoing bipolar division (**B**) and tripolar division (**C**). Scale bar = 10 um. Green dots = centrosomes (Cnn-GFP), Green cortical signal = membrane (Moesin-GFP), Magenta= DNA (his H2AV-RFP). Representative image from 5 animals, 34 cells, 2 replicates. (**D**) Timeline of applicable *Drosophila* papillar development at 25C along with a representative image of adult wildtype rectal papillae with *brachyenteron-(byn)-Gal4, Gal80^ts^*, *UAS-NLS* GFP. Dotted outlines = 1 papilla. Scale bar = 100um. (**E**) Left: Representative images of adult *UAS-shi RNAi* and *UAS-shi RNAi* + *plk4-OE* rectal papillae with *brachyenteron*-(*byn*)-*Gal4, Gal80^ts^*, *UAS-NLS* GFP. Dotted outlines = 1 papilla. Scale bar = 100um. Right: Graph of total number of rectal papillar cells in the adult fly with *UAS-shi* RNAi with and without *plk4-OE* as compared to wildtype (note that wildtype control experiment was done at a separate time from the *UAS-shi* RNAi experiments). Minimum n = 8 animals/group. p > 0.05 among all groups by unpaired T-test. Error bars represent mean and standard error. (**F**) EMS mutagenesis. (**F’**) Outline of the forward genetic screen. (**G**) Percent survival of all mutant lines (n = 642) with FLP/FRT-induced clones and *plk4-OE*-induced Centrosome Amplification (CA) after 10 days on a high salt diet. Red line indicates the 50% survival cut-ok used to identify the top 28 hits from the screen. Positive control = *Df(2R)Exel6062*, a chromosomal deficiency line that we previously found to repeatedly exhibit robust salt diet-dependent lethality (unpublished).

Compared to the study of centrosome clustering, very few studies have examined how cells may maximize survival following a complete multipolar division (**Figure 1A**, **outcome IIa vs IIb**). Hepatocytes of the mammalian liver routinely undergo CA and multipolar division, and such divisions lead to viable aneuploid daughter cells (Duncan et al., 2010, 2012; Guidotti et al., 2003). We previously reported that *Drosophila melanogaster* hindgut rectal papillar cells (hereafter: papillar cells) undergo CA and viable multipolar division that does not impact organ development or function (Schoenfelder et al., 2014). Both hepatocytes (Donne et al., 2020; Faggioli et al., 2011; Milne, 1909; Wang et al., 2017) and papillar cells are polyploid (Fox et al., 2010; Schoenfelder et al., 2014), meaning these cells have extra sets of the genome. Such extra genomes are likely crucial to surviving multipolar division, as extra copies of each chromosome per cell can buffer chromosome imbalances caused by aneuploidy (Ben-David & Amon, 2020). Approximately one-third of human cancers are polyploid (Bielski et al., 2018), and CA is highly prevalent in nearly all solid and hematological malignancies (Chan, 2011; Hu et al., 2022; Mittal et al., 2021). Currently, there is a gap in knowledge regarding molecular mechanisms that enable cells with CA to survive in the absence of centrosome clustering-i.e. to complete a viable multipolar cell division.

In this study, we exploited the error-prone nature of mitosis in *Drosophila* papillar cells (Bretscher & Fox, 2016; Clay et al., 2021, 2022; Fox et al., 2010; Schoenfelder et al., 2014; Stormo & Fox, 2016, 2019) to identify factors involved in the survival of cells with CA *in vivo*. Through a forward genetics screen of 642 chemically mutagenized fly lines, we identify the uncharacterized Drosophila gene *CG3168* as a novel vulnerability of cells with CA. Here, we name this gene *synaptic vesicle glycoprotein 2* (*sv2*) after the orthologous genes in mammals. We further show that a *Drosophila* SV2 transgenic protein localizes to the plasma membrane. In cells with CA, SV2 plays a critical role in centrosome positioning and chromosome congression during mitosis, allowing for higher fidelity multipolar divisions. While SV2 proteins have previously been examined in synaptic transmission and neural disease, this study reveals a novel role for SV2 in mitotic cells with CA.

## Results

### An *in vivo* forward genetic screen for novel vulnerabilities of cells with amplified centrosomes

We previously found that the papillar cells of the *Drosophila* hindgut undergo both bipolar divisions (**Figure 1B**, **Movie S1**) and multipolar divisions (**Figure 1C**, **Movie S2**) (Schoenfelder et al., 2014). These divisions take place during a 24-hour window of pupal development, when the adult gut is being formed (**Figure 1D-20-48 hours post-puparium formation (HPPF)**, **additional information in Figure S1A**). Despite the aneuploidy that results from multipolar division, papillar cells are incredibly tolerant of such divisions, which are viable. Additionally, papillar development and cell viability is remarkably unaffected by experimentally elevating levels of CA and multipolar division (Schoenfelder et al., 2014) through overexpression of *polo like kinase 4* (*plk4*-OE, also known in *Drosophila* as *SAK*), a kinase involved in centrosome duplication (Basto et al., 2008; Bettencourt-Dias et al., 2005; Cunha-Ferreira et al., 2009; Hudson et al., 2001; Rogers et al., 2009).

The high viability of papillar cells after multipolar division prompted us to investigate the underlying cellular survival mechanisms. We subsequently found that approximately 24 hours after the mitotic period, papillar tissue forms a giant syncytium (**Figure 1D-67-74 HPPF**) (Peterson et al., 2020). This process requires the GTPase dynamin (*Drosophila shibire-shi*). Syncytium formation could prevent lethal impacts of aneuploidy that result from multipolar divisions through cytoplasmic sharing of the products of imbalanced chromosomes. To test this hypothesis, we knocked down *shi* with and without *plk4*-OE-induced CA. There is no CA-dependent loss in the number of nuclei in the adult papillae (**Figure 1E-*shi RNAi* vs. *shi RNAi****, **plk4-OE***). These results suggest that syncytium formation is not a major factor in cell survival following multipolar division in papillar tissue.

Given the lack of a role for syncytium formation in tolerance of multipolar division-induced aneuploidy, we undertook a forward genetic screen to uncover novel vulnerabilities of cells with amplified centrosome number. We employed a strategy that enables rapid identification of EMS-induced mutations on the X chromosome (Haelterman et al., 2014). This strategy benefits from the ability to quickly map the candidate mutation to a small region of the X-chromosome by crossing to a library of BAC insertions containing X-chromosome rescue fragments (Cook et al., 2010; Erickson & Spana, 2006; Venken et al., 2010).

As with the previous X chromosome approach on which we based our screen (Haelterman et al., 2014), we aimed to induce mosaic FLP/FRT clones in our tissue of interest. Thus, our EMS mutagenesis was performed in an FRT19A genetic background that enables X chromosome mosaic clone generation. After recovering 6,617 mutagenized lines (methods), we identified those lines that were recessive lethal. For screening, we then crossed the recessive FRT19A lethal lines (**Figure 1F**, **mutant/+**) to a line containing an FRT19A chromosome and an inducible FLP recombinase (**Figure 1F’**, **FLP/FRT**). To optimize FLP recombinase induction, we experimented with an FRT19A-linked RFP reporter to determine an optimal FLP recombinase protocol that maximized papillar clone induction (methods). Our optimal protocol produced nearly 50% RFP+ and 50% RFP-papillar cells after clone induction (**Figure S1B**). Our previous work on the gene *fancd2* found that animals with roughly 50% papillar cell loss leads to an easily scoreable whole animal phenotype-death on a high salt diet (Bretscher & Fox, 2016). Therefore, we reasoned that any of our EMS-induced X chromosome mutations that are homozygous papillar cell lethal, either on its own or synthetic lethal with amplified centrosomes, would result in organismal lethality on a high salt diet.

To incorporate CA into our screen, our cross that introduced FLP recombinase also carried the hindgut-specific Gal4 driver *brachyenteron* (*byn*)-Gal4 and *UAS-SAK/plk4* (**Figure 1F’**, ***plk4*-OE**, methods). Therefore, resulting progeny where we induced mosaic lethal mutations also induced centrosome amplification in the developing hindgut, including papillar cells. Importantly, clone induction and CA occurred prior to any multipolar mitosis in papillar cells, which we previously showed to occur in the pupal phase (**Figure 1D**, **Figure S1A**) (Schoenfelder et al., 2014). Ultimately, our screen assayed for the ability of a recessive X chromosome mutation to cause a decrease in papillar cell number in the presence of extra centrosomes, with the readout being adult animal death with a high salt diet (**Figure 1F’**, **high salt diet**).

From our high salt diet screening of 642 FLP/FRT clones of X-chromosome recessive lethal lines, we identified 24 lines that repeatedly showed at least 50% animal lethality after 10 days (**Figure 1G**). These lines were carried forward for secondary screening.

In our secondary screen, we assessed whether mutant clone induction and CA were truly responsible for a given candidate identified from our primary screen. As with our primary screen, a positive control chromosomal deficiency stock that consistently results in organismal lethality on a high salt diet was included to ensure the sensitivity of our high salt assay (**Figure 2A**, **B**, **positive control**). Of the 24 lines identified from the primary screen, 6 lines ultimately became unhealthy without high salt treatment and were therefore excluded from further study. Therefore 18 lines that scored as hits in our primary screen were again tested for viability on high salt. In our first secondary screen, we omitted the induction of both mutant clones and CA. Removing these two primary screen criteria restored viability in all cases, underscoring the need to homozygose each recessive lethal mutation to observe a phenotype (**Figure 2A**). Similarly, for these same 18 lines, inducing clones without CA restores viability to above the 50% threshold (**Figure 2B**). We note that 3 lines were just barely above the threshold, suggesting these screen hits may be CA-independent (**Figure 2B**). These results confirm that a majority of the hits in our primary screen represent CA-dependent hits.

**Figure 2.**
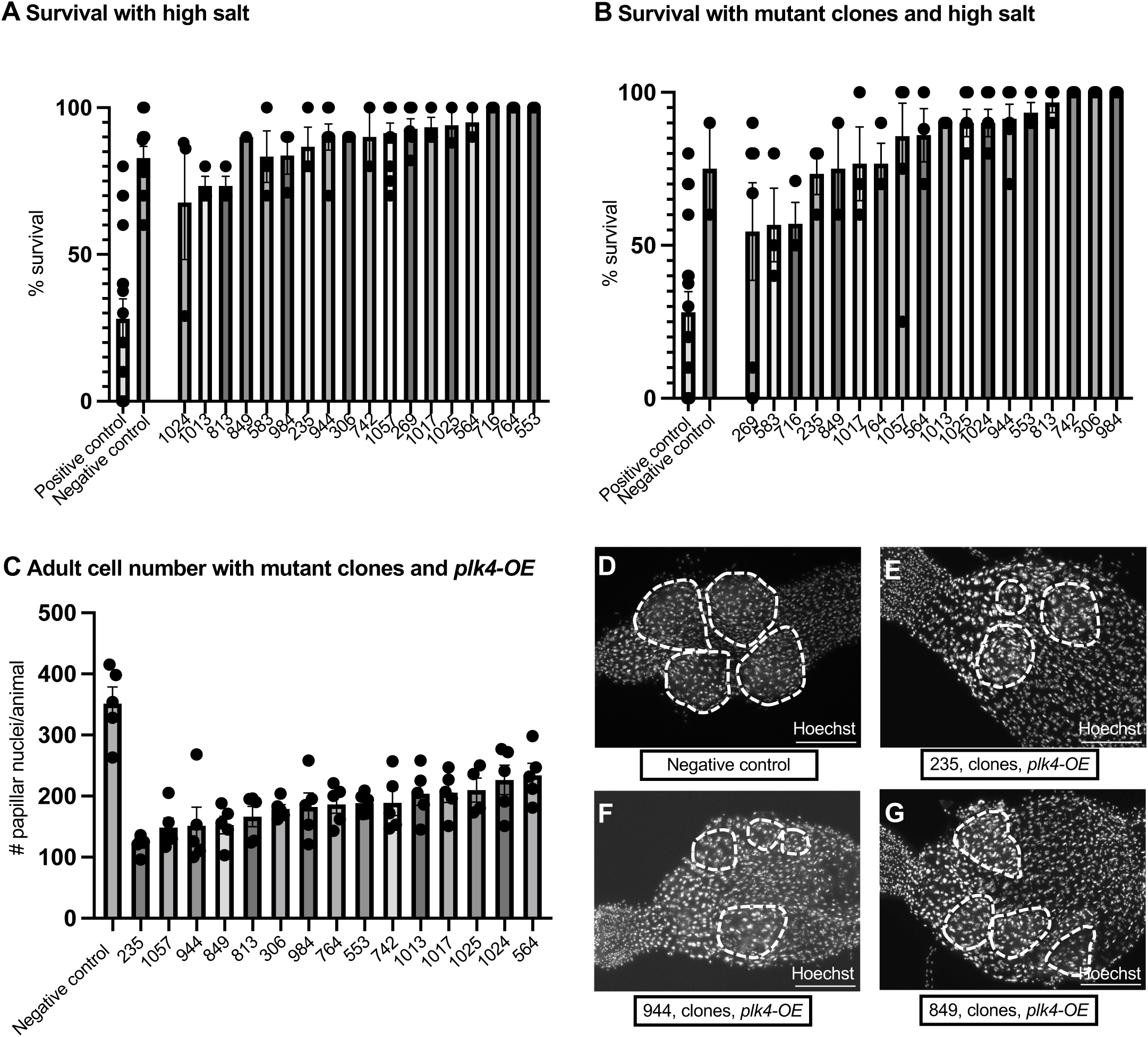
Secondary screening validates fifteen CA-dependent vulnerabilities. (**A**) Percent survival of X chromosome recessive lethal lines without *FLP/FRT 19A*-induced clones after 10 days on a high salt diet (minimum 10 animals/line). Positive control is *Df(2R)Exel6062*. Negative control is *FRT19A*. Error bars represent mean and standard error. (**B**) Percent survival of X chromosome recessive lethal lines with hs-FLP/FRT 19A-induced clones after 10 days on a high salt diet (minimum 9 animals/line). Positive control is *Df(2R)Exel6062*. Negative control 1 is *FRT19A; hs-FLP* with heat shock to induce clones. Error bars represent mean and standard error. (**C**) Number of nuclei in the adult papillae of the top 15 lines identified from the primary screen with *FLP/FRT 19A*-induced clones and *byn-Gal4, plk4-OE*-induced CA (minimum n=4 animals/line). Error bars represent mean and standard error. Negative control is *byn-gal4, UAS-GFP*. (**D-G**) Representative images of rectums from top hits with FLP/FRT 19A-induced clones and CA via *byn-Gal4* and *plk4-OE*. Dotted outlines = 1 papilla. Scale bar = 100um. (**D**) Negative control (*byn-gal4, UAS-GFP)*, (**E**) *FRT19A* 235, (**F**) *FRT19A* 944, (**G**) *FRT19A* 849.

As depicted in **Figure 1A**, we hypothesize that factors representing vulnerabilities of cells that frequently undergo multipolar division promote cell viability with CA. Given this, we tested if validated hits from our primary and secondary organism-level screens are required at the tissue level for achieving normal numbers of papillar cells with CA. We thus examined adult papillar tissue structure and cell number in animals where we induced CA and FLP/FRT clones of the X chromosomes from our validated 15 CA-dependent hits. Indeed, all tested mutants exhibit a sharp reduction in papillar cell number (**Figure 2C**), accompanied by visibly smaller or misshapen adult papillar organs as compared to control papillae (examples in **Figure 2D-G**). Ultimately, our forward genetic screen revealed 15 mutant lines that represent novel vulnerabilities of cells with CA in an *in vivo* context.

### *CG3168,* a major vulnerability of cells with centrosome amplification, encodes a ***Drosophila* Synaptic Vesicle Glycoprotein 2 family member**

In parallel with our secondary screening, we began to genetically map many of the validated screen hits to a region of the X chromosome by crossing EMS mutant lines to a collection of X chromosome fragment rescue lines (methods). One of the first mutants to be mapped was line 235, which has the most severe effect on papillar cell survival in the presence of FLP and *plk4*-OE (**Figure 2C**, **E**), while showing no appreciable effect on cell number in the absence of *plk4*-OE (**Figure S1C**). The organismal viability of recessive lethal mutant 235 is rescued by two X duplications (**Figure 3A**) that overlap by only 8,079 base pairs (**Figure 3B**). Within this small overlap region, there is only one gene coding sequence, which corresponds to the unnamed *CG3168* (**Figure 3B**). To test whether mutant 235 is an allele of *CG3168,* we generated a *CG3168* coding sequence transgene with a small FLAG-tag on the 5’ end, driven by a strong *ubiquitin* (*ubi)* promoter (**Figure 3C-FLAG-SV2**, methods). We performed crosses between heterozygous 235 females and heterozygous (single copy) ubi-FLAG-*CG3168* males and successfully recovered viable hemizygous 235 males (235/Y), albeit at sub-Mendelian ratios (**Figure 3D**, **1 transgene copy**). In subsequent crosses between 235/Y males containing a single copy of the transgene and 235 heterozygous females containing another single transgene copy, we successfully derived a healthy, viable, and fertile stock where the 235 mutant chromosome could be propagated as a homozygote along with two ubi-FLAG-*CG3168* copies (**Figure 3D**, **2 transgene copies**). Importantly, the FLAG-*CG3168* transgene fails to rescue organismal viability of the second strongest tissue-level screen hit, mutant 1057, speaking to the specificity of our *CG3168* rescue assay (**Figure 3D**). These results indicate dose-dependent rescue of strain 235 to full viability by a *CG3168* transgene, indicating that 235 is an allele of *CG3168*.

**Figure 3.**
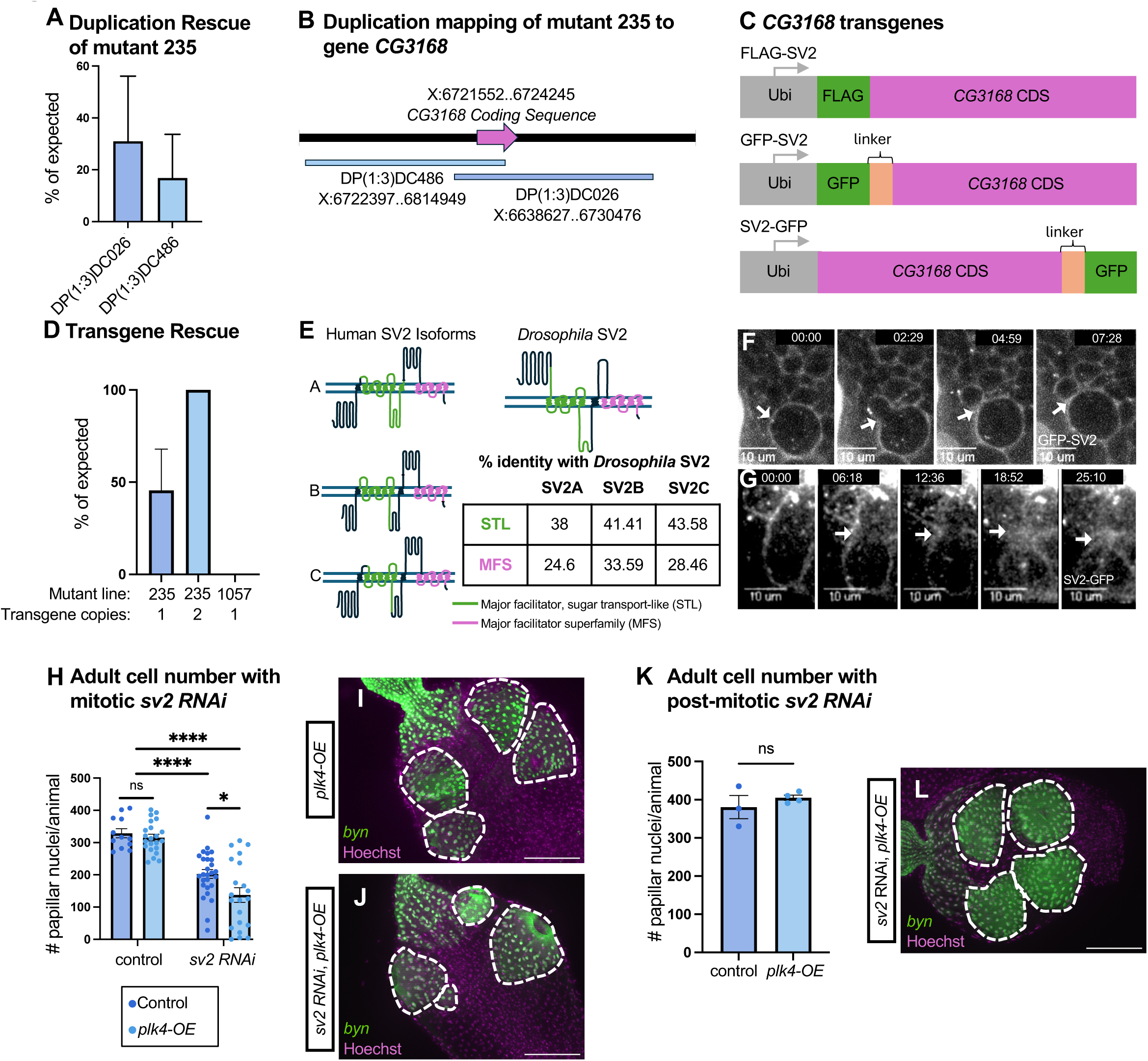
Identification and localization of SV2 as a gene of interest from the screen. (**A**) Organismal lethality rescue of mutant 235 with two duplication stocks (Cook et al., 2010) (minimum 20 animals/group). Error bars represent mean and standard error. (**B**) Mapping of the duplication stocks that rescue mutant 235 overlap on the coding sequence of *CG3168*. 5’ is to the left. (**C**) A schematic of *CG3168* transgene with a 5’ FLAG tag (top), 5’ GFP tag (middle) or 3’ GFP tag (bottom). 5’ is to the left. (**D**) Rescue level of mutant lethality with the indicated dose of FLAG-tagged *CG3168* transgene (minimum 38 animals/group). Error bars represent mean and standard error. (**E**) Protein structure comparison of the three isoforms of human SV2 and *Drosophila* CG3168, as well as percent identity comparison between human and *Drosophila* orthologs at the two conserved domains: the Major Facilitator Superfamily (MFS) and Major Facilitator, Sugar Transport-Like (STL) domains. Protein diagrams derived from Protter software predictions (methods). (**F**) 5’ GFP-tagged SV2 transgene expression during mitosis in a larval neuroblast. Scale bar = 10 um. Time measured in minutes with start at 0:00 being arbitrary at the start of the montage. Arrowheads indicate cleavage furrow in mitotic cell (**G**) 3’ GFP-tagged SV2 transgene expression during mitosis in a pupal rectal papillar cell. Scale bar = 10um. Time measured in minutes with start at 0:00 being arbitrary at the start of the montage. Arrowheads indicate cleavage furrow in mitotic cell. (**H**) Number of nuclei in the adult papillae with mitotic *UAS-sv2 RNAi^GD3258^* as compared to without *UAS-sv2 RNAi^GD325^*, with and without CA via *plk4-OE* at 27C (minimum n = 12 animals/group). Significance of *RNAi^GD325^* p<0.0001 and *plk4-OE* p = 0.0244 by Two-way ANOVA. Significant difference via unpaired T-test denoted on graph where * = p<0.05, ** = p<0.01, and *** = p<0.001. Error bars represent mean and standard error. (**I**) Representative image of a *UAS-plk4-OE*; *byn-Gal 4, Gal80^ts^, UAS NLS GFP* rectum with mitotic induction of *plk4-OE*. Dotted outlines = 1 papilla. Scale bar = 100um. (**J**) Representative image of a *UAS-plk4-OE; byn gal 4, gal80^ts^, UAS NLS GFP/UAS sv2 RNAi^GD3258^* rectum with mitotic induction of *sv2 RNAi^GD3258^* and *plk4-OE*. Dotted outlines = 1 papilla. Scale bar = 100um. (**K**) Number of nuclei in the adult papillae with post-mitotic *sv2 RNAi^GD3258^* with CA via *plk4-OE* as compared to without *plk4-OE* at 29C (minimum n = 3 animals/group from 1 replicate) p = 0.3988 via unpaired T-test. Error bars represent mean and standard error. (**L**) Representative image of a *UAS-plk4-OE; byn-Gal 4, Gal80^ts^, UAS NLS GFP/UAS sv2 RNAi^GD3258^* rectum with post-mitotic induction of *RNAi* and *plk4-OE*. Dotted outlines = 1 papilla. Scale bar = 100 um.

As *CG3168* is an unnamed gene, we examined homology of this top hit from our centrosome amplification vulnerability screen. *CG3168* encodes a protein with homology to solute carrier transmembrane proteins found throughout archaea, bacteria, and eukarya (Bajjalieh et al., 1992; Stout et al., 2019). In mammals, there are three proteins with high homology to the protein product of CG3168, named Synaptic Vesicle Glycoproteins (SV) SV2A, SV2B, and SV2C (**Figure 3E**). Mammalian SV2 proteins are best known for their role in vesicular transport and are most studied in neuronal signaling (Rossi et al., 2022; Stout et al., 2019). However, SV2 proteins are also expressed in various endocrine tissues outside the nervous system, as well as throughout the gastrointestinal tract (Portela-Gomes et al., 2000; Stout et al., 2019). From BLAST search data (Alliance of Genome Resources Consortium, 2024; Thurmond et al., 2019), *Drosophila* appear to have a single SV2 family member. As with mammalian SV2 family member expression, FlyAtlas2 RNAseq data (Krause et al., 2022; Leader et al., 2018) suggest that *CG3168* is widely expressed beyond the nervous system and therefore is likely to have functions beyond synaptic transmission. The protein sequence of SV2 is well conserved from humans to *Drosophila*, especially between the two major domains of the protein, the major facilitator superfamily domain (Jacobsson et al., 2007) and the major facilitator sugar transport-like domain (Madeo et al., 2014), as outlined in **Figure 3E** (also see Methods). We henceforth name the *CG3168* gene in *Drosophila sv2*, after the orthologous human gene(s).

The most prominent homology between fly and human SV2 proteins lies in predicted transmembrane domains (**Figure 3E**), and SV2 proteins are known to localize to the plasma membrane and associated vesicles (Feany et al., 1992; Lynch et al., 2004; Paulussen et al., 2024; Son et al., 2000; Xu & Bajjalieh, 2001). Therefore, we next examined the subcellular localization of GFP-tagged SV2 transgenic protein. We generated two Ubi-driven GFP transgenes, with the tag at either the 5’ or 3’ end of SV2, separated by a small linker sequence (**Figure 3C-GFP-SV2, SV2-GFP**). We focused on mitosis, given our finding of *sv2* as a hit in a CA vulnerability screen. We examined GFP-SV2 in mitotic neuroblasts. This transgenic protein strongly decorates the cortex of dividing neuroblasts, enabling us to observe asymmetric division and cytokinesis in live imaging (**Figure 3F**, **movie S3-arrow indicates cytokinesis**). In live mitotic stage papillar cells, we examined SV2-GFP. As with neuroblasts, we observe some cortical localization and can identify cytokinesis in papillar cells (**Figure 3G**, **movie S4-arrow indicates cytokinesis**), though the cortical GFP signal of this 3’ tagged SV2 transgenic protein in papillar cells is not as prominent as the signal of 5’ tagged SV2 in neuroblasts. Taken together, a top hit from our CA vulnerability screen corresponds to the single *Drosophila sv2* gene, the product of which may function at the plasma membrane during mitosis.

### CG3168/SV2 functions specifically during papillar cell mitosis

To further explore the function of SV2 in papillar cells, we examined two independent *sv2* RNAi lines (methods). We induced hindgut tissue-specific (via the *brachyenteron,* or *byn,* promoter) (Singer et al., 1996), temperature-inducible (via the *Gal4, Gal80^ts^* system) (McGuire et al., 2004) *UAS-sv2 RNAi* expression. Using this approach, we further assessed the role of *sv2* in papillar cells with CA.

Knockdown of *sv2* via both RNAi lines causes some decrease in number of papillar cells (**Figure 3H-J**, **Figure S2A)**. We did not observe this decrease when inducing clones of the 235 allele of *sv2* in the absence of CA (**Figure S1C**). This difference is most easily explained by the inability to induce more than 50% mutant cells of the 235 allele through FLP/FRT recombination, compared to 100% of papillar cells that can express the RNAi construct. Consistent with our findings with the 235 allele, *sv2* loss from either RNAi line sensitizes papillar cell number in the presence of CA (**Figure 3H-J**, **Figure S2A**). Along with our two duplication and FLAG-rescue gain of function assays, these loss of function results with two separate RNAi lines further support our identification of mutant 235 as an allele of *CG3168/sv2* in our screen for CA vulnerability.

We next assessed the window of development when *sv2* promotes survival of cells with CA. We previously defined the developmental events that produce rectal papillae (**Figure S1A**) (Fox et al., 2010; Peterson et al., 2020). To identify the critical period of SV2 function, we induced *sv2 RNAi* at distinct time points and examined the papillar cells at multiple developmental phases. First, to test whether SV2 is important for achieving normal cell numbers after completion of mitosis, we induced *sv2 RNAi* in post-mitotic stage pupae and examined papillar nuclear number in adults (methods). We find there is no significant decrease in papillar cell number from *sv2 RNAi* with or without CA after mitosis (**Figure 3K**-**L**, **Figure S2B**). We additionally find that SV2 is not necessary for cell survival prior to mitosis, as driving *sv2 RNAi* from embryogenesis through larval development has no impact on the number or appearance of rectal cells in wandering 3^rd^ instar larvae (methods, **Figure S2C-F**). The severe decrease of papillar cells only seen from *sv2* knockdown during mitosis suggests an important role for SV2 in the polyploid mitotic divisions of papillar cells, particularly upon CA.

### SV2 stabilizes centrosome positioning during mitosis

To uncover SV2 function during papillar cell mitosis with and without CA, we employed live imaging during the mitotic period (methods, **Figure S1A**). We compared mitosis in four separate genotypes: wildtype, *plk4-OE*, *sv2 RNAi*, and *sv2 RNAi + plk4-OE.* To maximize our ability to observe an enhancement of CA-related phenotypes by loss of SV2, we used a *plk4-OE* transgene insertion that leads to a moderate level (26%) of multipolar divisions in the absence of any RNAi. As previously described (Schoenfelder et al., 2014), papillar cells of all genotypes frequently exhibit CA and multipolar division, though some divisions exhibit clustering of extra centrosomes and bipolar division. The earliest difference among genotypes that we observe is that *sv2 RNAi* increases the length of metaphase in a CA-independent manner (**Figure 4A-D**, **Movie S5-7**). These results suggest a role of SV2 in regulation of the metaphase to anaphase transition.

**Figure 4.**
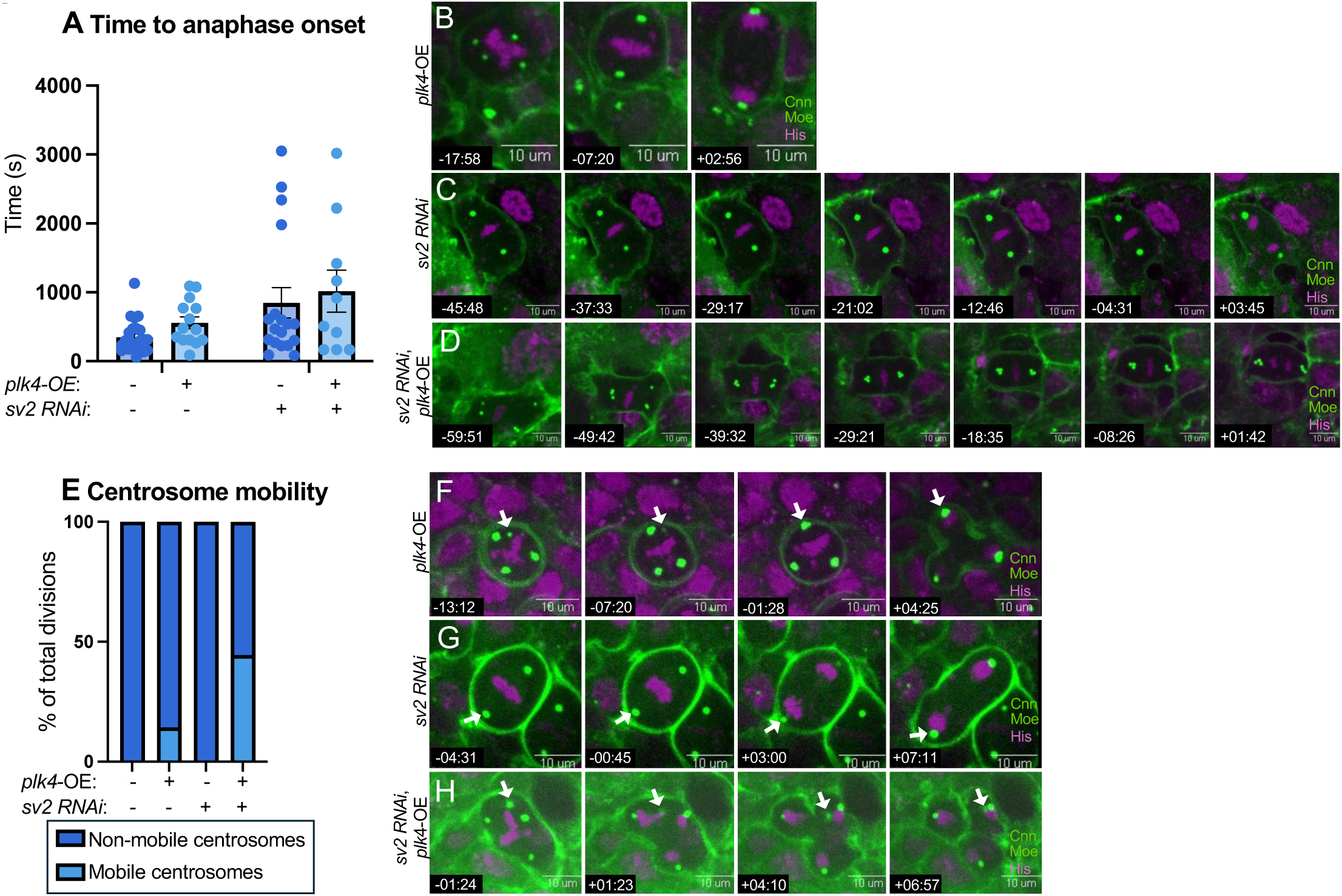
SV2 regulates anaphase timing and centrosome positioning during mitosis. (**A**) Time to anaphase onset in animals of the indicated genotypes. Minimum n= 10 cells from minimum 3 animals, minimum 2 replicates. *sv2 RNAi ^GD32^*. Error bars represent mean and standard error. The length of metaphase is significantly longer in *sv2 RNAi* compared among control and *plk4-OE* by one way ANOVA (P=0.0251). The length of metaphase is significantly longer in *sv2 RNAi + plk4-OE* compared among control and *plk4-OE* by one way ANOVA (P=0.0035). Time to anaphase measured as time from when metaphase plate was first visualized until the first time of chromosome separation. Time in measured in minutes relative to anaphase onset. (**B-D**) Representative montages from movies of the indicated genotypes. Green dots = centrosomes (*cnn-GFP*), Green cortical signal = membrane (*Moesin-GFP*), Magenta= DNA (*his H2AV-RFP*). *byn-Gal4* drives all transgenes except *H2AV-RFP,* which is driven by genomic sequence upstream of the gene. (**E**) Centrosome mobility in in animals of the indicated genotypes. Minimum n = 20 cells/genotype, n = 5 animals. (**F-H**) Representative montages from movies of the indicated genotypes. Green dots = centrosomes (*cnn-GFP*), Green cortical signal = membrane (*moesin-GFP*), Magenta= DNA (*his H2AV-RFP*). Arrowheads highlight the change (or lack thereof) in position of specific centrosomes in each montage. Time is measured in minutes relative to anaphase onset.

Prolonged metaphase is often triggered by problems in spindle biorientation (Basto et al., 2004; Gordon et al., 2001; Rieder et al., 1994) which can be especially problematic in cells with CA. Consistent with this idea, in live imaging of papillar cells, *sv2 RNAi* + *plk4-OE* cells frequently exhibit mobile centrosomes that appear unable to be stably anchored to the plasma membrane prior to anaphase onset (**Figure 4E-H**, **Movie S8-10**). Importantly, the increased centrosome mobility occurs after anaphase onset, not prior, when centrosomes are known to be mobile during processes such as spindle formation and centrosome clustering (Kwon et al., 2008; Marotta et al., 2025; Mercadante et al., 2023; Rhys et al., 2018; Yim et al., 2024) (**Figure 4F**). Our findings suggest that SV2 stabilizes centrosome position during mitosis, and that this stability is especially challenged upon CA. As a result, *sv2 RNAi* + *plk4-OE* papillar cells frequently prolong metaphase and enter anaphase with an unstable multipolar centrosome configuration.

Following the initial period of increased centrosome mobility, *sv2 RNAi + plk4-OE* papillar cells exhibit additional catastrophic mitotic properties related to metaphase chromosome alignment. Impaired chromosome congression to the metaphase plate can also cause anaphase delay and can be complicated by merotelic kinetochore attachments (attachments where at least one sister chromatid kinetochore remains engaged with more than one spindle pole) (Cimini et al., 2001, 2003; Janicke & LaFountain, 1982; Nicklas & Arana, 1992; Rieder et al., 1995). Merotelic attachments are more common in cells with CA (Drpic et al., 2018; Ganem et al., 2009; Silkworth & Cimini, 2012). Consistent with the idea that *sv2* loss may cause alignment defects in a CA-dependent manner, *sv2 RNAi + plk4-OE* papillar cells have numerous unaligned chromosomes that fail to congress to the metaphase plate (**Figure 5A-D**, **Movie S11-13**). Additionally, loss of *sv2* enhances the frequency of multipolar division, regardless of whether we experimentally elevate CA by *plk4-OE*, further suggesting a role for *sv2* in promoting viability of cells with CA (**Figure 5E-H**, **Movie S14-16**). To summarize, we find that SV2 promotes efficient and timely progression from metaphase to anaphase. In papillar cells with CA, SV2 ensures stable centrosome position and chromosome congression. Overall, our findings here establish at the organismal, tissue, and single cell level a critical and novel role for an SV2 family member in mitotic fidelity, especially in cells that exhibit CA and multipolar division.

**Figure 5.**
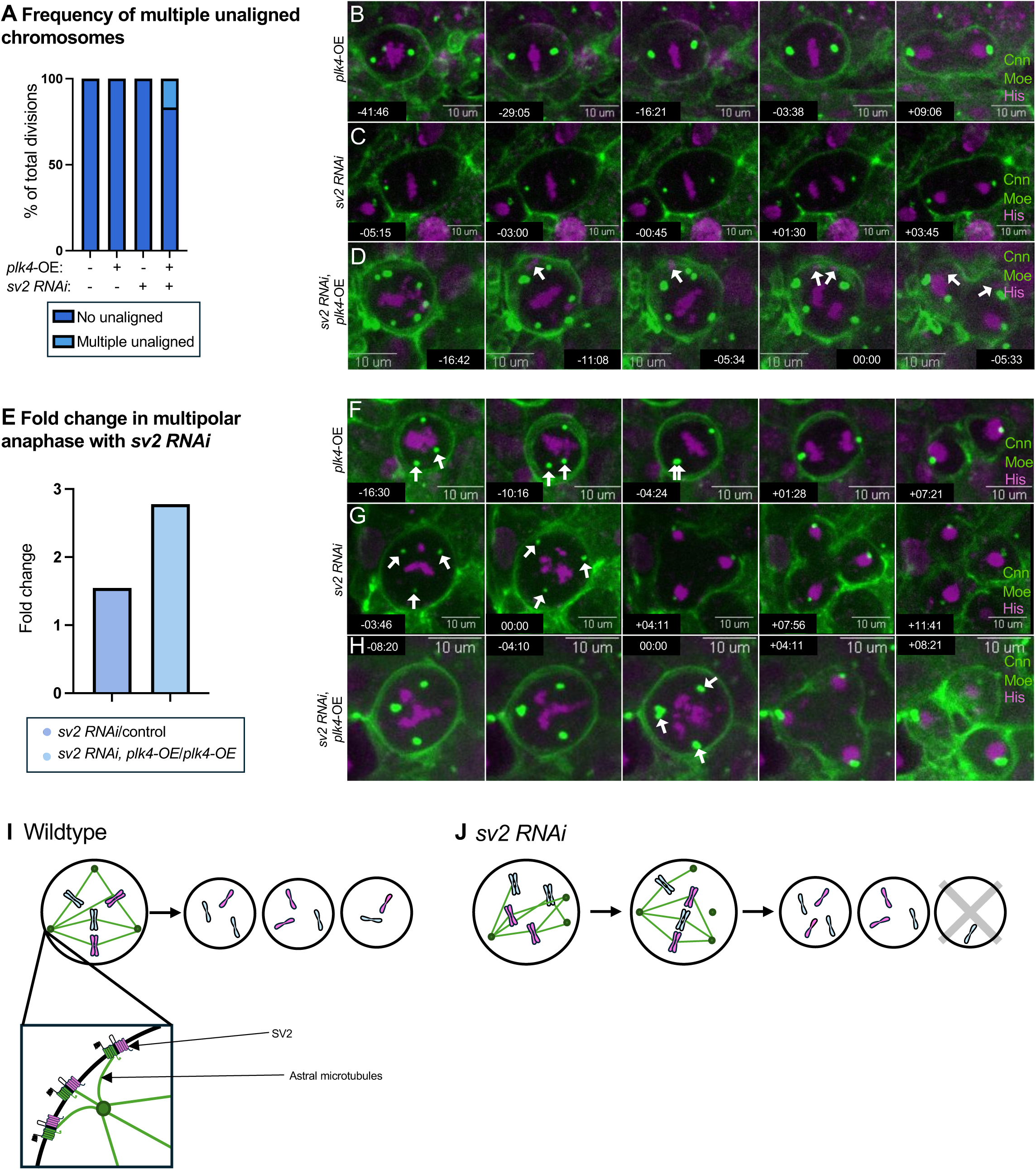
SV2 regulates metaphase congression and division outcome in cells with CA. (**A**) Frequency of multiple unaligned chromosomes at metaphase in animals of the indicated genotypes. Minimum n = 18 cells, n = 4 animals. (**B-D**) Representative montages from movies of the indicated genotypes. Green dots = centrosomes (cnn-GFP), Green cortical signal = membrane (*moesin-GFP*), Magenta= DNA (*his H2AV-RFP*). *byn-Gal4* drives all transgenes except *H2AV-RFP,* which is driven by genomic sequence upstream of the gene. Arrowheads indicate unaligned chromosomes. Time is measured in minutes relative to anaphase onset. (**E**) Fold change of multipolar division with *UAS-sv2 RNAi^GD32^* as compared to cells without *sv2 RNAi^GD32^*, both with and without *plk4*-OE. Ratio of *sv2 RNAi^GD32^*/control = 1.55. Ratio of *plk4-OE* + *sv2* RNAi*^GD32^*/*plk4-OE* = 2.78. (**F-H**) Representative montages from movies of the indicated genotypes. Green dots = centrosomes (*cnn-GFP*), Green cortical signal = membrane (*moesin-GFP*), Magenta= DNA (*his H2AV-RFP*). Arrowheads highlight centrosomes. Time is measured in minutes relative to anaphase onset. (**I**) Model for SV2 function during multipolar division. (**J**) Model for multipolar mitosis in the absence of SV2.

## Discussion

In this study, we performed an unbiased forward genetic screen to identify a previously unknown factor for the survival of cells with extra centrosomes. Our X chromosome recessive lethal screen and follow-up secondary screens ultimately found 15 distinct lines where we demonstrated clear CA-dependent decrease in adult cell number. In addition to reporting our screen results, this study characterizes our first studied CA-dependent hit. We name this factor SV2, due to orthology to human SV2 family proteins for this previously unnamed *Drosophila* protein. SV2 acts during the mitotic period to achieve normal numbers of cells in the papillar tissue, and, like the mammalian ortholog, an SV2 transgenic protein localizes to the plasma membrane in multiple cell types. Finally, through live imaging of papillar cell mitosis, we reveal a role for SV2 in multiple aspects of mitosis. Some of these functions (lengthened anaphase) are apparent without CA, which suggest a fundamental role for SV2 in mitotic fidelity. Additional SV2 functions are found in cells with CA, underscoring why we identified an allele of SV2 as a top hit in our screen and follow-up analysis.

### A Model for the Role of SV2 in the Survival of Cells with Extra Centrosomes

Together, our data suggest that plasma membrane-localized SV2 coordinates interactions of centrosomes with the mitotic spindle. Our data lead us to suggest that SV2 interacts with the astral microtubules of the spindle, ensuring the fidelity of centrosome placement and stability at the spindle poles during metaphase and prior to anaphase onset (**Figure 5I**). Without SV2, we speculate that there is a corresponding loss of the integrity of mitotic spindle attachments, thereby increasing time to anaphase onset via activation of the spindle assembly checkpoint, as well as increasing centrosome mobility. This increase in mobility is especially challenged in cells with CA and likely causes the multiple unaligned chromosomes that we observe in the absence of SV2 in cells with CA (**Figure 5J**).

Interaction of SV2 with the mitotic spindle would also explain the problems with chromosome congression to the metaphase plate that we observe in cells with increased centrosome number. CA places an undue strain on the cell to perform mitosis with high fidelity (Ganem et al., 2009; Silkworth & Cimini, 2012). We propose that this decrease in mitotic fidelity leads to cell death after division, which ultimately decreases the number of adult papillar cells. Future work, beyond the scope of this genetic screen and initial characterization, can refine our model by examining spindle structure regulation by SV2 in detail. Our findings here suggest the importance of membrane integrity in the fidelity of cell division *in vivo*, in line with the idea that chromosome segregation relies upon correct tissue architecture (Knouse et al., 2018).

### A Novel Role for Synaptic Vesicle Glycoprotein 2 Proteins in Cell Division

The SV2 family of proteins in mammals has been largely studied in the brain, with a focus on its role in neurotransmission (Ciruelas et al., 2019; Stout et al., 2019). In this study, we describe a previously unknown role for an SV2 ortholog in *Drosophila* as a regulator of mitotic fidelity, especially in cells with amplified centrosomes. *Drosophila* papillar cells are a model of how evolution can adapt to tolerate error-prone mitosis and the resultant aneuploidy (Fox et al., 2020). This study provides new mechanistic insight into this process. Given the conserved nature of SV2 proteins, it is of paramount importance to next explore the role of SV2 proteins in cell division in other organisms.

Papillar cells have proven to be an applicable model for studying cancer-relevant biology (Bretscher & Fox, 2016; Clay et al., 2021, 2022). Our work here suggests that targeting SV2 proteins (for which there is an FDA-approved drug, Levetiracetam, that targets human SV2A) (Gower et al., 1992; Lynch et al., 2004) may be a viable mechanism to suppress growth of tumor cells that tolerate multipolar cell division (Aroosa et al., 2023). Indeed, the SV2 family of proteins in humans are expressed in several different types of cancer, including pancreatic, brain, lung, male and female reproductive, liver, colorectal, and urothelial (Karlsson et al., 2021; Uhlén et al., 2005). More recently, pharmacological inhibition of SV2A has been shown to inhibit the growth of murine neuroendocrine prostate cancer cells (Sulsenti et al., 2021). Future study can examine whether SV2 inhibition is particularly effective in tumors with polyploidy and multipolar division.

In summary, our work here highlights the continued power of unbiased *in vivo* genetic screens to reveal unexpected roles for conserved proteins. It is amazing that cells have mechanisms that enable them to tolerate splitting into more than two daughter cells and living to tell the tale. We look forward to continued study in this fascinating and disease-relevant area of biology.

## Materials and methods

### Drosophila culturing

Flies were kept on standard *Drosophila* medium (Archon scientific) except where high salt food was used (see “salt diet feeding” below). Transgenic flies generated for this study were produced by Jamie Roebuck (*Drosophila* genetic services):

*ubi-FLAG-CG3168*

*ubi-GFP-CG3168*

*ubi-CG3168-GFP*

### EMS mutant generation

Adult males from an isogenized FRT19A stock were fed 10mM EMS. Resulting mutagenized X chromosomes were placed over an FM7 balancer chromosome and individual lines were established. Lines were then screened for recessive whole organism lethal mutations, revealed by the absence of males without an FM7 chromosome. Of 6,617 initial lines, we obtained 1,500 lethal lines, of which we ended up screening 642. We reasoned that lethal lines were most valuable for screening, as they likely contained an important molecular lesion.

### FLP-FRT clone induction

We determined our optimal heat-shock protocol by separately generating *FRT19A, ubi-RFP+* marked homozygous clones with *hs-FLP12*, and identifying the developmental timing and heat-shock duration that achieved ∼50% clones, the theoretical maximum. Importantly, we visualized clones prior to papillar tissue syncytium formation (Peterson et al., 2020), after which individual clones are no longer visible.

### EMS screen and salt diet feeding

*hs-FLP12, FRT19A;; byn-Gal4, UAS-SAK / TM3, Sb* males were crossed to *FRT19A* EMS mutant females, each of which were over an *FM7* balancer. To induce homozygous clones, the resulting progeny were heat-shocked twice for 40 min. upon embryo hatching/early L1 stage, to maximize clones in hindgut papillar precursor cells. Resulting female non-*Sb* progeny were placed on food containing 250mM NaCl for ten days and then scored for viability.

### Genetic mapping, transgene rescue, and complementation

Crosses between EMS mutant females and males containing either an X chromosome duplication or a transgene of interest were scored for the presence of viable F1 males with the lethal mutation-containing chromosome that were now viable as hemizygotes, indicating rescue. Duplication lines contain Bacterial Artificial Chromosomes with fragments of the X chromosome on autosomes or are the product of a translocation between the X and Y chromosome that resulted in a duplication (Cook et al., 2010; Erickson & Spana, 2006; Venken et al., 2010). For most lines, a clear single region of the X chromosome was identified as a candidate region of interest. For transgene rescue, Twist biosciences cloned the cDNA for a candidate gene downstream of a ubiquitin promoter and *Drosophila* injection services generated transgenic cDNA rescue animals, which we then crossed back to the original parental line and assayed for rescue of viability, as was done for the X chromosome mapping.

### SV2 Domain Structure

STL and MFS domains of SV2 orthologs were identified using UniProt and FlyBase. Percent identity of SV2 human orthologs to Drosophila SV2 were identified using BLASTp alignment. Protein diagrams adapted from transmembrane protein prediction server, Protter (https://wlab.ethz.ch/protter/start/)(Omasits et al., 2014).

### RNAi

For inducing *UAS-RNAi* constructs prior to mitosis and during endocycling for larval dissections, crosses were set between the RNAi line and *byn gal4*, *gal80^ts^ UAS NLS GFP +/-UAS-SAK (plk4-OE)*. Egg laying occurred for 48 hours at 18C. The vials were then shifted from 18C to 29C at 72 hours to induce *UAS-RNAi*. Third instar larvae from the cross were then picked from vials to be dissected and analyzed. For *UAS-RNAi* induction during mitosis, the same laying conditions were used, and vials were similarly shifted from 18C at 72 hours to either 29C or 27C. The eclosed adults of those crosses were then dissected and analyzed. For post-mitotic induction of *UAS-RNAi*, the same laying conditions were used, but the vials were shifted from 18C to 29C on day 16 (approximately half-way through pupariation, after mitosis). The eclosed adults were then dissected and analyzed. Only females were analyzed for both adult *UAS-sv2 RNAi* lines to control for any sex-specific variables.

### Fixed imaging

All tissues were dissected and fixed as previously described (Fox et al., 2010; Fox & Spradling, 2009). All fixed imaging was performed using a Zeiss AxioImager M.2 with Apotome processing at 20X magnification with 1um z slice sections. All fixed images were analyzed using FIJI using Z projection and cell counter tools.

### Live imaging

All tissues were dissected and processed for live imaging as previously described (Fox et al., 2010; Fox & Spradling, 2009) and imaged using an Andor XD Spinning Disk Confocal Microscope on 60X magnification with a silicon oil objective. All live imaging analysis and movie assembly was performed using MetaMorph software. Time to anaphase onset was measured as time from the first visualization of the metaphase plate until first visualization of chromosome separation at anaphase. Movie timestamps are in minutes relative to anaphase onset. There is an exception for figures 3F and G, where movie timestamps are in minutes and 0:00 is arbitrary for the first image of the montage.

## Acknowledgments

The authors are deeply grateful for discussions between DF and the late Dr. Angelika Amon, who encouraged us to perform this EMS screen. The following kindly provided reagents used in this study: Bloomington Drosophila Stock Center, Vienna Drosophila Resource Center. The Duke Light Microscopy Core Provided imaging support. We thank Drs. Scott Hawley (FlyBoard Nomenclature Committee), Kevin Cook (Bloomington Drosophila Stock Center), and Steven Marygold (FlyBase) for discussions on gene naming. We thank Drs. Terry Lechler, Corinne Linardic, Masayuki Onishi, and Lee Zou (Duke University) and Fox lab members for comments on the manuscript. This project was supported by NIH grant R01 GM140138 and NASA Translational Institute for Space Health (TRISH) grant NNX16AO69A-T0108 to DF.

