## Supplemental Material for "Synaptic vesicle glycoprotein 2 enables viable aneuploidy following centrosome amplification"

### Figure S1

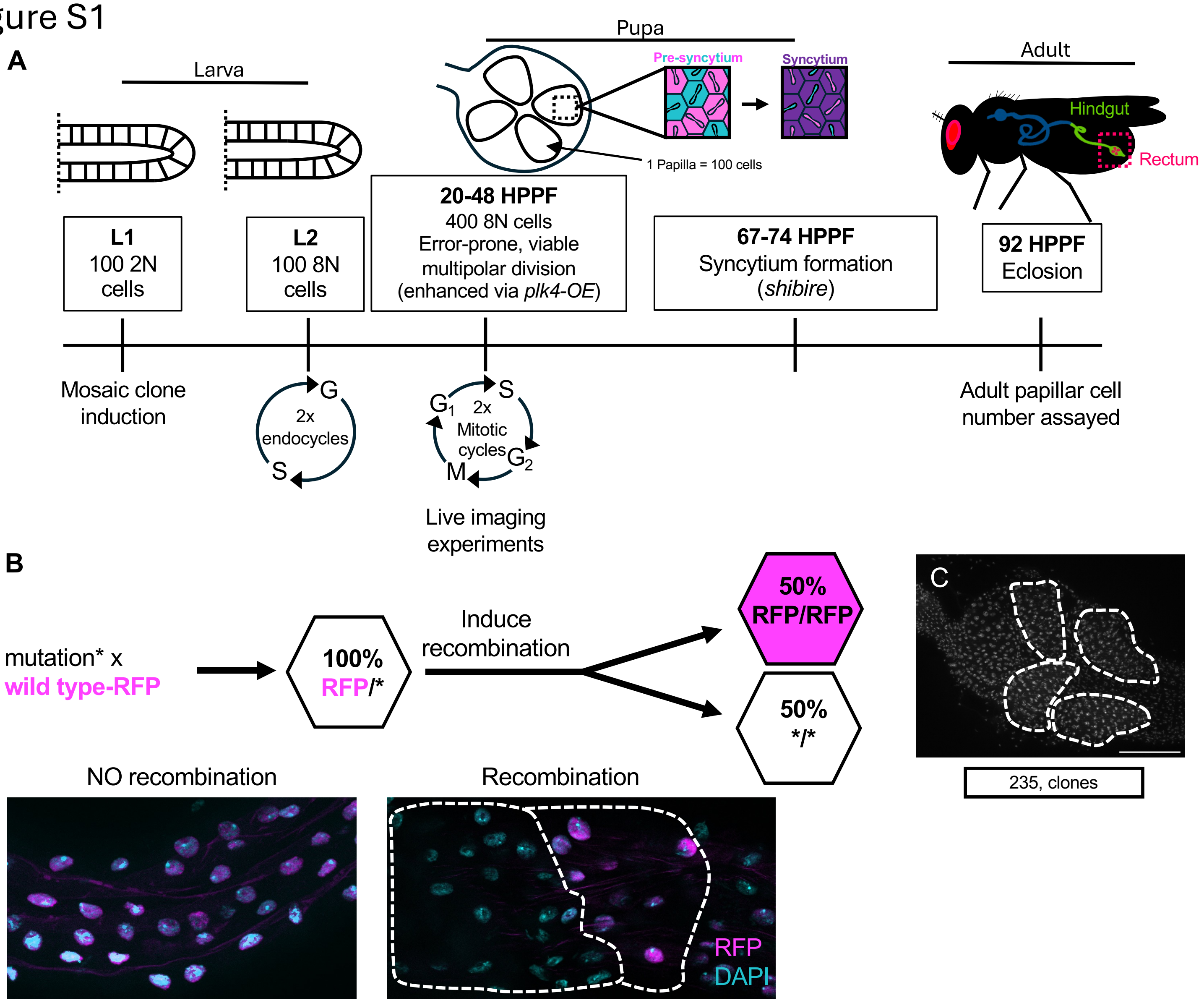
A
Larva
Pupa
Adult
Pre-syncytium
Syncytium
Hindgut
Rectum
1 Papilla = 100 cells
L1
100 2N cells
L2
100 8N cells
20-48 HPPF
400 8N cells
Error-prone, viable multipolar division
(enhanced via plk4-OE)
67-74 HPPF
Syncytium formation
(shibire)
92 HPPF
Eclosion
Mosaic clone induction
G
2x endocycles
S
S
G1
2x Mitotic cycles
G2
M
Live imaging experiments
Adult papillar cell number assayed
B
mutation* x
wild type-RFP
100%
RFP/*
Induce recombination
50%
RFP/RFP
50%
*/*
C
235, clones
NO recombination
Recombination
RFP
DAPI

Figure S2

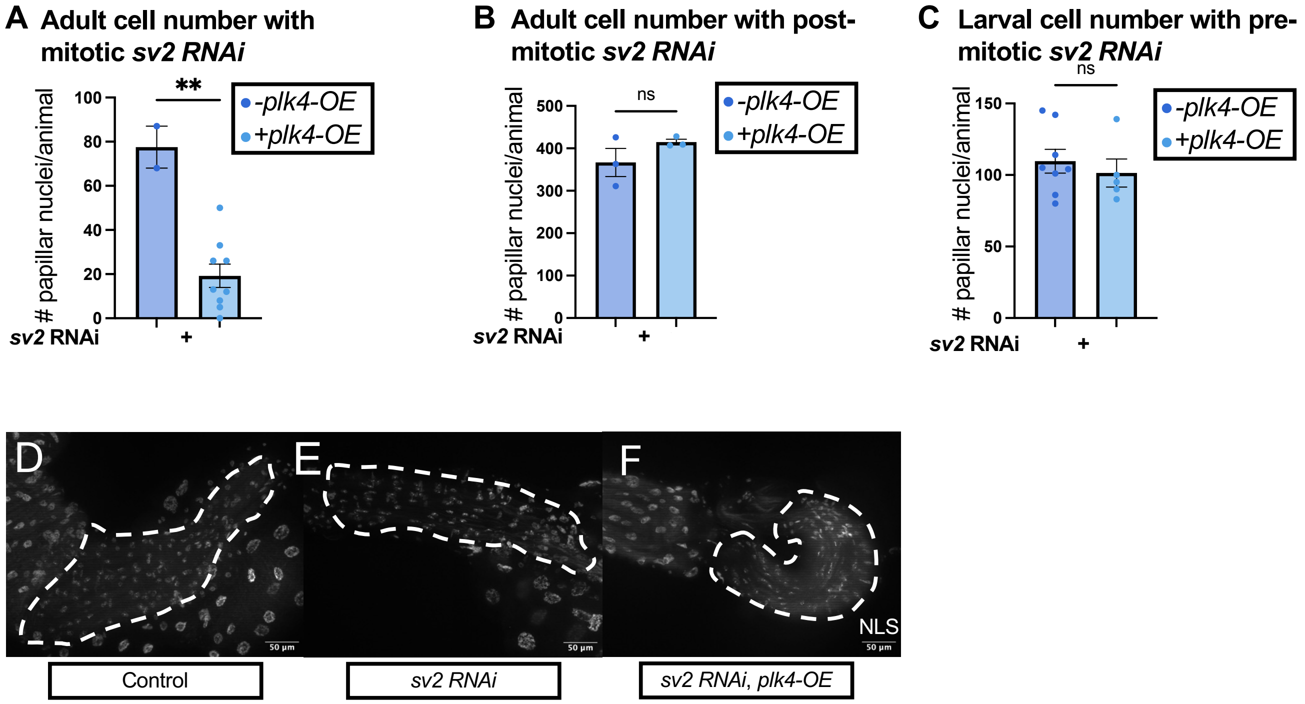

A Adult cell number with mitotic sv2 RNAi
**
-plk4-OE
+plk4-OE
### papillar nuclei/animal
100
80
60
40
20
0
sv2 RNAi
+
B Adult cell number with post-mitotic sv2 RNAi
ns
-plk4-OE
+plk4-OE
### papillar nuclei/animal
500
400
300
200
100
0
sv2 RNAi
+
C Larval cell number with pre-mitotic sv2 RNAi
ns
-plk4-OE
+plk4-OE
### papillar nuclei/animal
150
100
50
0
sv2 RNAi
+
D
E
F
NLS
50 µm
50 µm
50 µm
Control
sv2 RNAi
sv2 RNAi, plk4-OE

**Supplemental Material**

**2 Supplemental Figures**

**16 Supplemental Movies**

**Supplemental Figure Legends**

**Figure S1. (A)** Timeline of *Drosophila* papillar development at 25C from first larval instar stage through adulthood. **(B)** Experimental diagram of induction of clones in hsflp12, *Frt19A/Frt19A, Ubi-RFP (NLS);; byn gal4, UAS plk4-OE, UAS FLP/+* rectum via heat shock as well as representative images of non-heat shock control and heat-shocked experimental rectum with twin spots (dotted lines). (**C**) Representative image of a 235 mutant line rectum with FLP/FRT 19A-induced clones without CA. Dotted outlines = 1 papilla. Scalebar = 100 um.

**Figure S2. (A)** Number of cells in the adult papillae with mitotic *UAS-sv2 RNAi*$^{\mathrm{TRiP.JF02441}}$ with and without CA via *plk4-OE* at 29C (minimum n = 2 animals/group from 1 replicates). p= 0.0010 via unpaired T-test. **(B)** Number of cells in the adult papillae with post-mitotic *sv2 RNAi* $^{\mathrm{TRiP.JF02441}}$ with and without CA via *plk4-OE* at 29C (minimum n = 3 animals/group from 1 replicate) p = 0.2299 via unpaired T-test. **(C)** Number of cells in the adult papillae with pre-mitotic *sv2 RNAi* $^{\mathrm{TRiP.JF02441}}$ with and without CA via *plk4-OE* at 29C (minimum n = 5 animals/group from 1 replicate) p = 0.5429 via unpaired T-test. **(A-C)** Error bars represent mean and standard error. **(D-F)** Representative images of WL3 rectums with wildtype phenotype **(D)**, with *UAS-sv2 RNAi*$^{TRiP.JF02441}$ **(E)**, or with *UAS-sv2 RNAi*$^{TRiP.JF02441}$ and *plk4-OE* **(F)**.

Dotted lines = 1 larval rectum. Scale bar = 50um (**D-F**).

**Supplemental Movies**

**Supplemental Movie 1. Bipolar division in a wildtype papillar cell.**

**Supplemental Movie 2. Tripolar division in a wildtype papillar cell.**

**Supplemental Movie 3. GFP-tagged SV2 in a mitotic larval brain neuroblast.**

**Supplemental Movie 4. GFP-tagged SV2 in a mitotic pupal papillar cell.**

**Supplemental Movie 5. Mitosis duration in a *plk4-OE* papillar cell.**

**Supplemental Movie 6. Mitosis duration in an *SV2 RNAi* papillar cell.**

**Supplemental Movie 7. Mitosis duration in an *SV2 RNAi, plk4-OE* papillar cell.**

**Supplemental Movie 8. Centrosome positioning in a *plk4-OE* papillar cell.**

**Supplemental Movie 9. Centrosome positioning in an *SV2 RNAi* papillar cell.**

**Supplemental Movie 10. Centrosome positioning in an *SV2 RNAi, plk4-OE* papillar cell.**

**Supplemental Movie 11. Chromosome alignment in a *plk4-OE* papillar cell.**

**Supplemental Movie 12. Chromosome alignment in an *SV2 RNAi* papillar cell.**

**Supplemental Movie 13. Chromosome alignment in an *SV2 RNAi, plk4-OE* papillar cell.**

**Supplemental Movie 14. Bipolar mitosis in a *plk4-OE* papillar cell.**

**Supplemental Movie 15. Multipolar mitosis in an *SV2 RNAi* papillar cell.**

**Supplemental Movie 16. Multipolar mitosis in an *SV2 RNAi, plk4-OE* papillar cell.**
